# The intersection of inflammation and DNA damage as a novel axis underlying the pathogenesis of autism spectrum disorders

**DOI:** 10.1101/2024.12.11.627854

**Authors:** Megha Jhanji, Colleen L. Krall, Alexis Guevara, Brian Yoon, Mathew Sajish, Luigi Boccuto, Sofia B. Lizarraga

## Abstract

Autism spectrum disorders (ASD) affects 1 in 36 children and is characterized by repetitive behaviors and difficulties in social interactions and social communication. The etiology of ASD is extremely heterogeneous, with a large number of ASD cases that are of unknown or complex etiology, which suggests the potential contribution of epigenetic risk factors. In particular, epidemiological and animal model studies suggest that inflammation during pregnancy could lead to an increased risk of ASD in the offspring. However, the molecular mechanisms that contribute to ASD pathogenesis in relation to maternal inflammation during pregnancy in humans are underexplored. Several pro-inflammatory cytokines have been associated with increased autistic-like behaviors in animal models of maternal immune activation, including IL-17A. Using a combination of ASD patient lymphocytes and stem cell-derived human neurons exposed to IL-17A we discovered a shared molecular signature that highlights a metabolic and translational node that could lead to altered neuronal excitability. Further, our work on human neurons brings forward the possibility that defects in the DNA damage response could be underlying the effect of IL-17A on human excitatory neurons, linking exacerbated unrepaired DNA damage to the pathogenicity of maternal inflammation in connection to ASD.

## INTRODUCTION

Large epidemiological studies suggest that maternal infection during pregnancy could be a major environmental risk factor for autism spectrum disorders (ASD) (Lee *et al*, 2015; Lydholm *et al*, 2019). Severe viral infections during pregnancy are associated with a one-third increase in ASD risk in offspring (Lee *et al*., 2015). For example, an analysis of 96,000 Danish children showed that risk for ASD doubled in response to severe maternal influenza infection (Atladottir *et al*, 2012). Evidence from clinical and postmortem studies suggests an emergent immune signature associated with ASD. Blood samples from individuals with ASD showed higher gene expression of pro-inflammatory genes such as TNF-α, IL-6, and IL-17 compared to controls (Eftekharian *et al*, 2018). Similarly, transcriptome analyses of postmortem brain tissues identified an overrepresentation of genes involved in immune regulation in ASD brains compared to unaffected individuals (Sciara *et al*, 2020; Voineagu *et al*, 2011). Work from animal models of maternal immune activation (MIA) suggests that a major mechanism underlying the effect of MIA during mammalian brain development is the IL-6/IL-17A immunoregulatory axis. MIA leads to elevated levels of IL-6, driving the differentiation of Th0 cells into Th-17 cells in the mother (Smith *et al*, 2007). These activated Th17 cells produce IL-17A (Kimura & Kishimoto, 2010), which can cross the placental barrier and disrupt fetal brain development. The direct effect of IL-17A in the developing cerebral cortex was shown in animal models; for example, in-utero injection of IL-17A into the embryonic mouse cortices at day 14.5 (E14.5), led to anatomical and behavioral phenotypes that resemble those associated with MIA (Choi *et al*, 2016). However, mechanistic studies on the role of IL-17A on cortical human neurons and how that might intersect with underlying mechanisms are lacking.

Transcriptome studies of rat fetal brains exposed to MIA during a developmental stage equivalent to the human gestational third trimester showed upregulation of pathways implicated in translation and DNA damage (Lombardo *et al*, 2018). In contrast, treatment of human forebrain organoids with activated IL-6 showed downregulation of genes implicated in translation (Sarieva *et al*, 2023) and implicated a primary effect on neuronal progenitor cells, suggesting a species and cell type-specific response to inflammatory signals in the developing brain. Emerging evidence highlights the therapeutic potential of resveratrol (RSV), a polyphenolic compound with anti-inflammatory and antioxidant properties, in counteracting neuroinflammation and oxidative stress, which are prominent features in ASD models. RSV treatment in animal studies has shown promise in alleviating ASD-like behaviors such as social deficits and repetitive behaviors (Bakheet *et al*, 2016; Bambini-Junior *et al*, 2014; Bhandari & Kuhad, 2017; Fontes-Dutra *et al*, 2018; Xie *et al*, 2018). However, recent findings suggest that RSV’s biological effects may differ between its *cis-* and *trans*-isomers, which have distinct mechanisms of action. For instance, these isomers differentially regulate the nuclear translocation and activity of tyrosyl-tRNA synthetase (TyrRS), a key enzyme involved in DNA repair and stress response (Cao *et al*, 2017; Jhanji *et al*, 2022). These differences could have significant implications for the therapeutic application of RSV in ASD, underscoring the need for isomer-specific studies.

Additionally, clinical evidence suggests that innate immune dysfunction in individuals with ASD may exacerbate behavioral symptoms (Hughes *et al*, 2023). This highlights the potential for therapeutic interventions targeting immune and oxidative stress pathways to extend benefits beyond the prenatal period. However, the extent to which there are common mechanisms altered by inflammation at fetal and postnatal stages in humans in relation to ASD pathogenesis is underexplored. Here, we use IL-17A as well as anti- and pro-inflammatory *cis* and *trans*-resveratrol (*cis*-RSV and *trans*-RSV) to dissect their effect on maturing human neurons mechanistically. We find that exposure to inflammatory signals impairs neurite outgrowth and decreases neuronal activity in stem cell-derived excitatory neurons. These cellular phenotypes correlate with alteration on key downstream signaling pathways, including (STAT3, MAPK, and NF-κB), as well as increased unrepaired DNA damage, which elicits the activation of the innate immune cGAS pathway (Liu *et al*, 2018) and leads to increased nuclear translocation of TyrRS where it activates DNA repair mechanisms (Cao *et al*., 2017). Additional transcriptomic and WGCNA analysis on stem cell-derived human neurons showed that inflammation leads to functional enrichment primarily on metabolic and translation pathways. Finally, we find that the key molecular signatures identified in stem cell-derived human neurons transcend cell types and genetic background as analysis of lymphocytes obtained from ASD individuals showed similar transcriptome changes.

## RESULTS

### IL-17A impairs neuronal morphogenesis and activity, with exacerbation by *trans*-RSV

ASD is a disorder of neuronal connectivity, and the proper development of neuronal connections requires precise regulation of neuronal arborization and synaptic function. To explore the impact of inflammatory signaling on human neurons, we used human embryonic stem cell (hESC) line H9 (Thomson *et al*, 1998) and differentiated hESCs into cortical excitatory neurons using the dual-SMAD inhibition protocol (Shi *et al*, 2012). We examined the effects of IL-17A, *cis*- and *trans*-RSV alone and in combination with IL-17A after 72 hours of treatment at day 55 of differentiation in cortical excitatory neurons (**Fig. 1A-E and Supplementary Fig. S1**). Neurite outgrowth analysis showed a significant decrease in mean and total neurite length with IL-17A and with IL-17A + *trans*-RSV but not with *cis*-RSV (**Fig. 1B-C**). Similarly, neuronal arborization, as assessed by complexity index, was reduced in IL-17A and IL-17A + *trans*-RSV conditions (**Fig. 1D**), while Scholl analysis had a similar non-significant trend for IL-17A alone (**Fig. 1E**). These findings indicate that IL-17A disrupts neuronal morphogenesis, with *trans*-RSV amplifying these effects.

**Figure 1.**
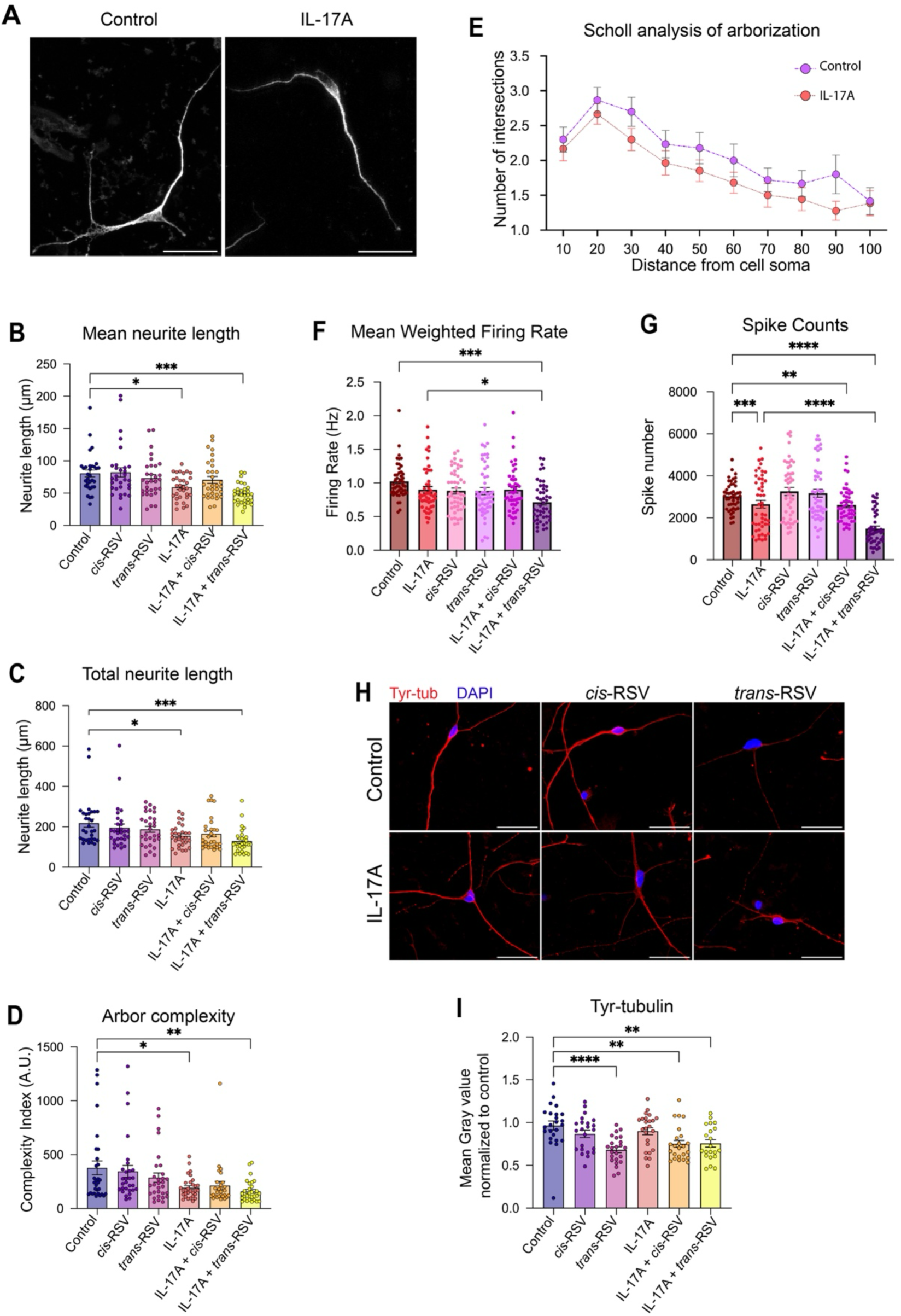
Effect of IL-17A, *cis* and *trans*-RSV on neuronal arborization, activity and tubulin tyrosination. (**A**) Representative images show hESC-derived neurons stained with MAP2 in black and white for ease of viewing. Scale bar is 30 µm. (**B-E**) Analysis of neuronal morphogenesis for control neurons (blue bars and circles), and neurons treated with either *cis-*Resveratrol (*cis-*RSV) (purple bars and circles), *trans-*Resveratrol (*trans-*RSV) (lilac bars and circles), IL-17A (red bars and circles), IL-17A + *cis-*RSV (orange bars and circles), and IL-17A + *trans-*RSV (yellow bars and circles). Mean and standard error of the mean (SEM) are shown for at least 30 neurons measured per condition. (**B**) Graph shows analysis of the mean neurite length; (**C**) Graph shows analysis of total neurite length; and (**D**) Graph shows analysis of arbor complexity by complexity index. Statistical analysis for (B-D) was done using One-way ANOVA with Dunnett’s test for multiple comparisons; *P <0.05, **P < 0.005, ***P <0.001. (**E**) Graph shows Scholl analysis for control (purple circles) and IL-17A (red circles) treated neurons. Statistical analysis by mixed model, P values that were not significant are not shown. (**F-G**) Analysis of neuronal activity by weighted mean firing rate (**F**) and spike counts (**G**) for control neurons (brown bar and circles), neurons treated with either IL-17A (red bar and circles), *cis-*RSV (pink bars and circles), *trans-*RSV (light purple bars and circles), IL-17A + *cis-*RSV (magenta bars and circles) and IL-17A + *trans-*RSV (dark purple bars and circles). The average of 3 independent experiments are shown with at least 8 wells per condition. The individual points represent the average per well across 12 different time points and across the 3 independent experiments per condition. Statistical analysis was conducted using grouped analysis by Two-way ANOVA test. *P <0.05, **P < 0.005, ***P <0.001, and ****P < 0.0001. (**H-I**) Analysis of tyrosinated tubulin levels in response to different treatments. (**H**) Representative images are shown stained with a tyrosinated tubulin antibody (red) and nuclei are stained with DAPI (blue). Scale bar is 30µm. (**I**) Quantification of mean gray values for tyrosinated tubulin was analyzed as mean gray value per field for control neurons (blue bars and circles), and neurons treated with either *cis-*RSV (purple bars and circles), *trans-*RSV (lilac bars and circles), IL-17A (red bars and circles), IL-17A + *cis-*RSV (orange bars and circles), and IL-17A + *trans-*RSV (yellow bars and circles). Mean ± SEM are shown with individual points representing individual fields. Statistical analysis was conducted using One-way ANOVA test, **P < 0.005, and ****P < 0.0001.

Next, we evaluated the effect of IL-17A, *cis*- and *trans*-RSV on neuronal activity using multi-electrode array (MEA) approaches after a 24-hour treatment for day 55 cortical excitatory neurons. Neurons for each treatment were cultured in 8 wells containing electrodes, and spontaneous neuronal activity was measured for 15 minutes every 2 hours for a total of 24 hours. By day 55 of neuronal induction cortical excitatory neurons are becoming synaptically active (**Fig. 1F-G**). Weighted mean firing rates showed a non-significant decrease in IL-17A-treated neurons but a significant reduction in neurons treated with IL-17A + *trans*-RSV (**Fig. 1F**). Spike counts were significantly lower with IL-17A alone and with IL-17A combined with either *cis-*RSV or *trans-*RSV, while no significant effects were observed with RSV isomers alone (**Fig. 1G**). These results suggest that IL-17A reduces neuronal activity, with differential contributions from RSV isomers.

To dissect the mechanisms underlying these deficits, we examined dynamic microtubules, critical for neurite growth and synaptic plasticity, using tyrosinated tubulin as a marker (Hosseini *et al*, 2022; Moutin *et al*, 2021). Imaging analysis of cellular tyrosinated tubulin after 72 hours showed a reduction in tyrosinated tubulin in neurons treated with *trans-*RSV, IL-17A + *cis-*RSV, and IL-17A + *trans-*RSV, but not in IL-17A alone or *cis-*RSV alone (**Fig. 1H-I**). Western blot analysis after 4 hours of treatment showed a significant decrease in all conditions, with the largest decrease in neurons treated with IL-17A, *trans-*RSV, and combination treatments (**Supplementary Figure S1**). Reduced tyrosinated tubulin levels may reflect altered activity of tyrosine tubulin ligase (TTL), TyrRS, or both. Interestingly, total TTL levels increased in neurons treated with IL-17A, *trans-*RSV, IL-17A + *cis-*RSV and IL-17A + *trans-*RSV, potentially reflecting a compensatory response to preserve microtubule integrity. Total TyrRS levels also trend upward, achieving significance in IL-17A-treated neurons compared to control neurons (**Supplementary Figure S1**). However, imaging analysis after 4 hours of treatment showed reduced TyrRS levels in the nucleus, implying a nuclear depletion of TyrRS by IL-17A (**Supplementary Figure S1**). Together, these results highlight how inflammation and RSV isomers affect pathways regulating microtubule dynamics and neuronal homeostasis. In summary, IL-17A impairs neuronal morphogenesis and activity in human cortical neurons, with *trans*-RSV exacerbating these effects.

### IL-17A elicits activation of specific downstream signaling pathways in human neurons

The observed changes in tyrosinated tubulin suggest disruptions in microtubule dynamics, potentially driven by signaling pathways such as STAT3 and NF-κB (Jackman *et al*, 2009; Verma *et al*, 2009), both implicated in immune downstream signaling. In non-neuronal systems, IL-17A activates different signaling pathways, including NF-κB, JAK/STAT, and MAPK (Rex *et al*, 2023), while in human neuronal progenitor cells (NPCs), IL-17A stimulation leads to the activation of ERK, RPS6, and NF-κB signaling pathways (Gomes *et al*, 2022). To investigate these pathways in differentiated neurons, we examined day 55 cortical excitatory neurons following 4 hours of treatment (**Fig. 2A-F**). Analysis of total cell lysates showed that IL-17A, *trans-*RSV and IL-17A + *trans-*RSV all increased total STAT3 levels, but only IL-17A + *trans-*RSV significantly elevated phospho-STAT3 (p-STAT3) levels (**Fig. 2A-C**). In contrast, phospho-MAPK (p-MAPK) and NF-κB2 were significantly upregulated by IL-17A or *trans-*RSV individually, but not in combination (**Fig. 2D-E**). However, the total levels of NF-κB were upregulated with both *cis-* and *trans-*RSV as well as IL-17A + *trans-*RSV, while IL-17A alone had no significant effect (**Fig. 2F**). To clarify this discrepancy, we examined the nuclear localization of NF-κB as a marker of its activation. Nuclear NF-κB levels were significantly increased in neurons treated with IL-17A, *trans-*RSV, and IL-17A + *trans-*RSV but remained unchanged with *cis-*RSV treatment (**Fig. 2G-H**). This differential activation pattern suggests that IL-17A and RSV isomers engage distinct regulatory mechanisms influencing NF-κB signaling. Taken together, our biochemical and imaging analysis suggest that inflammatory signals activate key pathways, including STAT3, MAPK, and NF-κB, which may contribute to observed defects in microtubule dynamics.

**Figure 2.**
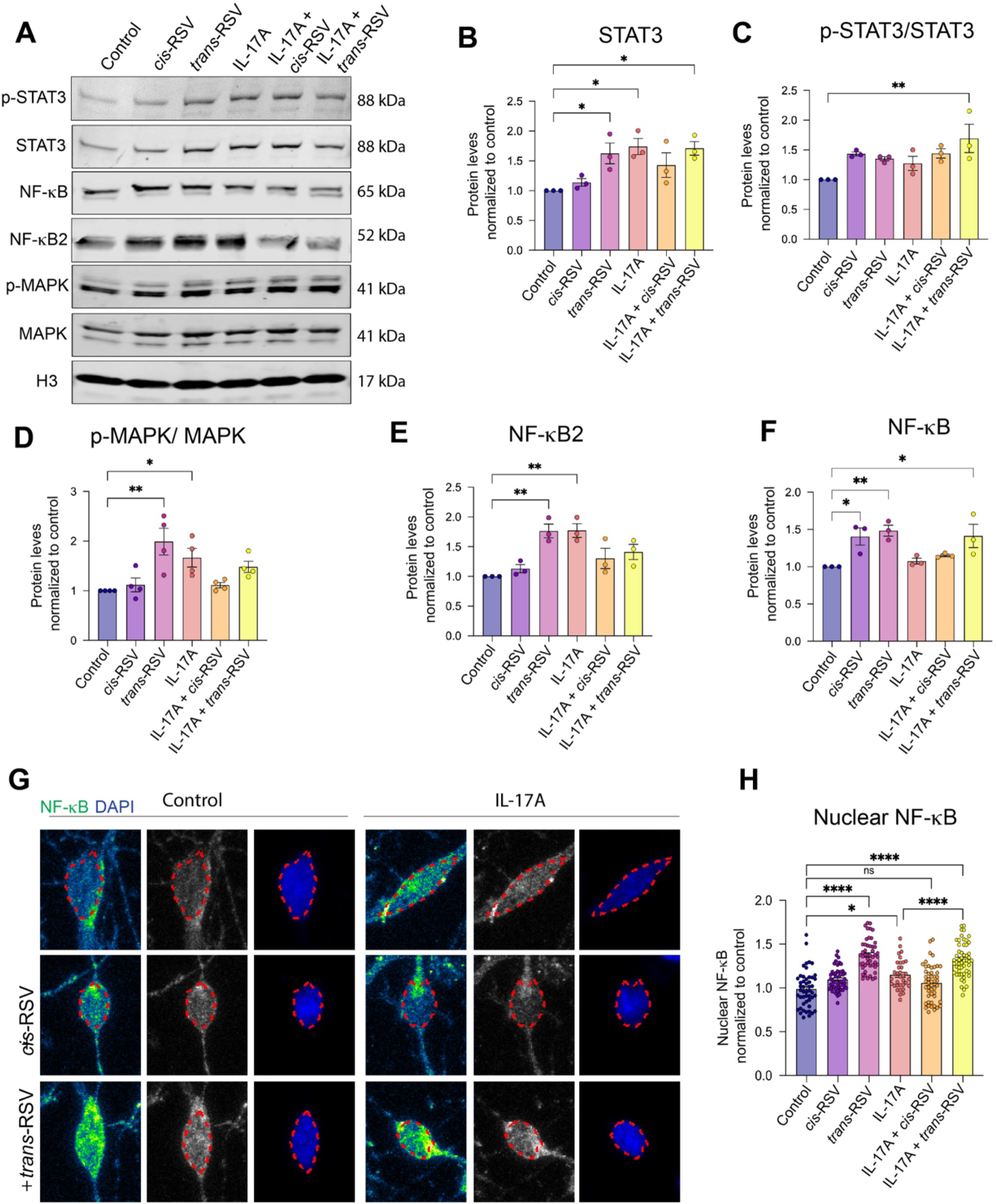
Analysis of downstream signaling pathways in human neurons. (**A**) Representative images show western blot for neuronal cell lysates obtained from either control neurons or neurons treated with IL-17A, *cis-*RSV, *trans-*RSV or IL-17A + *cis-*RSV and IL-17A + *trans-*RSV. Histone H3 was used as a loading control for total protein levels unless otherwise specified. Specific protein and their corresponding molecular weights are shown. (**B-F**) Analysis of protein levels for control neurons (blue bars and circles), and neurons treated with either *cis-*RSV (purple bars and circles), *trans-*RSV (lilac bars and circles), IL-17A (red bars and circles), IL-17A + *cis-*RSV (orange bars and circles), and IL-17A + *trans-*RSV (yellow bars and circles). Mean ± SEM are shown for at least three independent experiments for STAT3 (**B**), phospho-STAT3 (p-STAT3) normalized to STAT3 (**C**), phospho-MAPK normalized to MAPK (**D**), NF-κB2 (**E**) and NF-κB (**F**). Statistical analysis for (**B-F**) was done using ordinary One-way ANOVA; *P <0.05, and **P < 0.005. (**G-H**) Analysis of nuclear translocation of NF-κB in response to different treatments. (**G**) Representative images for neurons stained with an NF-κB antibody shown for which the signal is shown as the LUT (green) and nuclei are stained with DAPI (blue). Scale bar is 10µm. (**H**) Quantification of mean gray values for nuclear NF-κB was analyzed for 30 neurons in 3 independent experiments for control neurons (blue bars and circles), and neurons treated with either *cis-*RSV (purple bars and circles), *trans-*RSV (lilac bars and circles), IL-17A (red bars and circles), IL-17A + *cis-*RSV (orange bars and circles), and IL-17A + *trans-*RSV (yellow bars and circles). Mean ± SEM are shown with individual points representing the average across three independent experiments. Mean gray values were normalized to the control. Statistical analysis was conducted as grouped analysis using two-way ANOVA with Sidak’s test, *P < 0.05, and ****P < 0.0001.

### Accumulation of DNA damage as a response to inflammatory signals in human neurons

STAT3 and MAPK activation are well-documented contributors to DNA repair mechanisms in non-neuronal cells (Barry *et al*, 2010; Wood *et al*, 2009). Similarly, NF-κB activation has been linked to the cellular response towards DNA damage (Stilmann *et al*, 2009; Wang *et al*, 2017). However, in neurons, unresolved DNA damage can exacerbate inflammatory signaling by releasing nuclear DNA fragments into the cytoplasm, activating the cGAS-STING pathway (Li & Chen, 2018). Subsequent translocation of cGAS to the nucleus would further prevent DNA repair, amplifying genomic instability (Liu *et al*., 2018). We found that both *trans-*RSV and IL-17A increased nuclear cGAS levels (**Fig. 3A-B**), consistent with potential cytoplasmic DNA release and heightened immune activation. To directly assess DNA damage, we measured nuclear foci for phospho-histone H2A.X (γ-H2A.X), a marker of double-strand breaks (**Fig. 3C-D**). Neurons treated with IL-17A, *trans-*RSV or their combination showed significantly elevated γ-H2A.X foci, indicating increased accumulation of unrepaired DNA damage.

**Figure 3.**
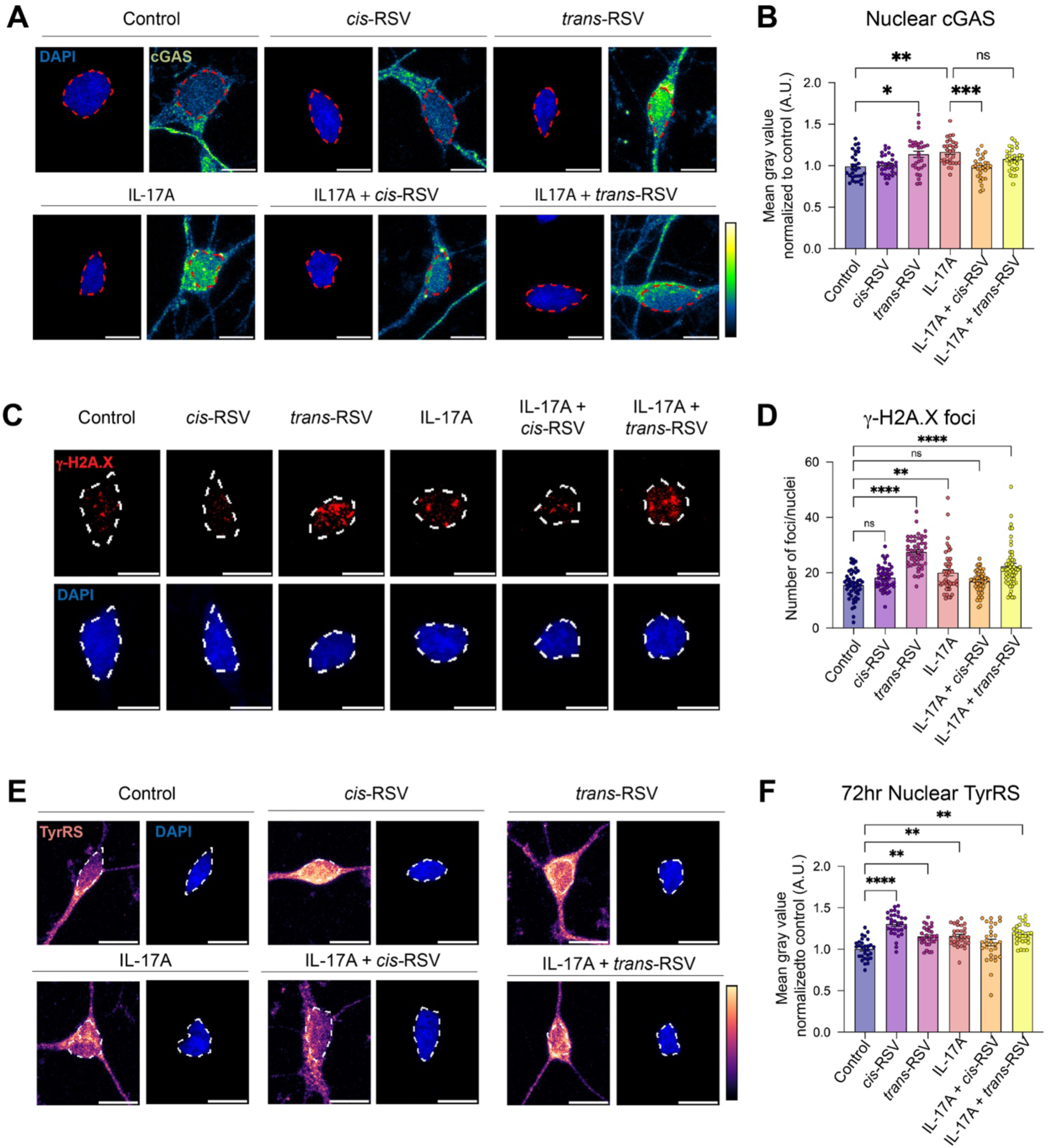
Analysis of the effect of inflammation on DNA damage in human neurons. (**A**) cGAS nuclear translocation in response to different treatments. Representative images of neurons stained with cGAS antibody are shown as the LUT (green) and nuclei are stained with DAPI (blue). Red outline shows the region that was measured. Scale bar is 10µm. (**B**) Quantification of mean gray values for nuclear cGAS was analyzed for 30 neurons in 3 independent experiments for control neurons (blue bars and circles), and neurons treated with either *cis*-Resveratrol (*cis-*RSV) (purple bars and circles), *trans*-Resveratrol (*trans*-RSV) (lilac bars and circles), IL-17A (red bars and circles), IL-17A + *cis-*RSV (orange bars and circles), and IL-17A + *trans*-RSV (yellow bars and circles). Mean ± SEM are shown with individual points representing the average across 30 neurons in three independent experiments. Mean gray values were normalized to the control. Statistical analysis was conducted as grouped analysis using two-way ANOVA with Sidak’s test for multiple comparisons, *P < 0.05, **P < 0.004, and ***P < 0.0009. (**C**) Analysis of γ-H2A.X nuclear foci in response to different treatments. Representative images of neurons stained with γ-H2A.X antibody (red) and nuclei stained with DAPI (blue) are shown. White outline shows the region measured. Scale bar is 10µm. (**D**) Quantification of the number of nuclear γ-H2A.X foci was analyzed for 30 neurons in 3 independent experiments for control neurons (blue bars and circles), and neurons treated with either *cis*-Resveratrol (*cis-*RSV) (purple bars and circles), *trans*-Resveratrol (*trans*-RSV) (lilac bars and circles), IL-17A (red bars and circles), IL-17A + *cis*-RSV (orange bars and circles), and IL-17A + *trans-*RSV (yellow bars and circles). Mean ± SEM are shown with individual points representing the average across 30 neurons in three independent experiments. Statistical analysis was conducted using one-way ANOVA with Sidak’s test for multiple comparisons, **P < 0.002 and ***P < 0.00000009. (**E**) TyrRS nuclear translocation in response to different treatments. Representative images of neurons stained with TyrRS antibody are shown as the LUT (pink hues) and nuclei are stained with DAPI (blue). White outline shows the region measured. Scale bar is 10µm. (**F**) Quantification of mean gray values for nuclear TyrRS was analyzed for 30 neurons in 3 independent experiments for control neurons (blue bars and circles), and neurons treated with either *cis*-Resveratrol (*cis-*RSV) (purple bars and circles), *trans*-Resveratrol (*trans*-RSV) (lilac bars and circles), IL-17A (red bars and circles), IL-17A + *cis*-RSV (orange bars and circles), and IL-17A + *trans*-RSV (yellow bars and circles). Mean ± SEM are shown with individual points representing the average across 30 neurons in three independent experiments. Mean gray values were normalized to control. Statistical analysis was conducted as grouped analysis using two-way ANOVA with Sidak’s test for multiple comparisons, **P < 0.0099, and ***P < 0.000000005.

Given the observed increase in total TyrRS levels in the IL-17A-treated neurons, we examined its nuclear translocation (**Fig. 2G**), which is known to promote DNA repair under oxidative stress (Wei *et al*, 2014). After a 72-hour treatment, we found nuclear TyrRS levels to be elevated in neurons treated with IL-17A, *cis-*RSV, *trans-*RSV, or IL-17A + *trans*-RSV (**Fig. 3E-F**). This finding suggests that inflammatory treatment may induce oxidative stress, prompting a compensatory response involving TyrRS nuclear localization to mitigate the DNA damage. Together, these findings demonstrate that IL-17A and the *trans-*RSV isomer exacerbate DNA damage and trigger stress response pathways, while *cis-*RSV appears to mitigate some of these effects, suggesting a potential protective role under inflammatory conditions.

### Pro-inflammatory cytokine IL-17A alters translation, mitochondrial and metabolic pathways in human neurons

To explore the molecular mechanisms disrupted by IL-17A in human neurons in relation to *cis-*RSV and *trans-*RSV, we analyzed transcriptomic changes at day 55 following a 24hrs treatment when neurons began to display neuronal activity (**Fig. 4, Supplementary Table S1 and Supplementary Fig. S2**). We identified a distinct set of differentially expressed genes (DEGs) in response to IL-17A alone (73 DEGs, FDR<0.1) (**Fig. 4A and Supplementary Fig. S2**). However, treatment with IL-17A + *cis-*RSV (143 DEGs, FDR <0.05) and IL-17A + *trans-*RSV (1044 DEGs, FDR<0.05) showed a more pronounced effect when compared to controls. Specifically, IL-17A + *trans-*RSV led to a significant upregulation of gene expression with 87.8% of DEGs showing significant overexpression (**Fig. 4B-C and Supplementary Fig. S2**). Combination treatments with RSV isomers showed molecular signatures that were distinct from each other (**Fig. 4D and Supplementary Fig. S2**).

**Figure 4.**
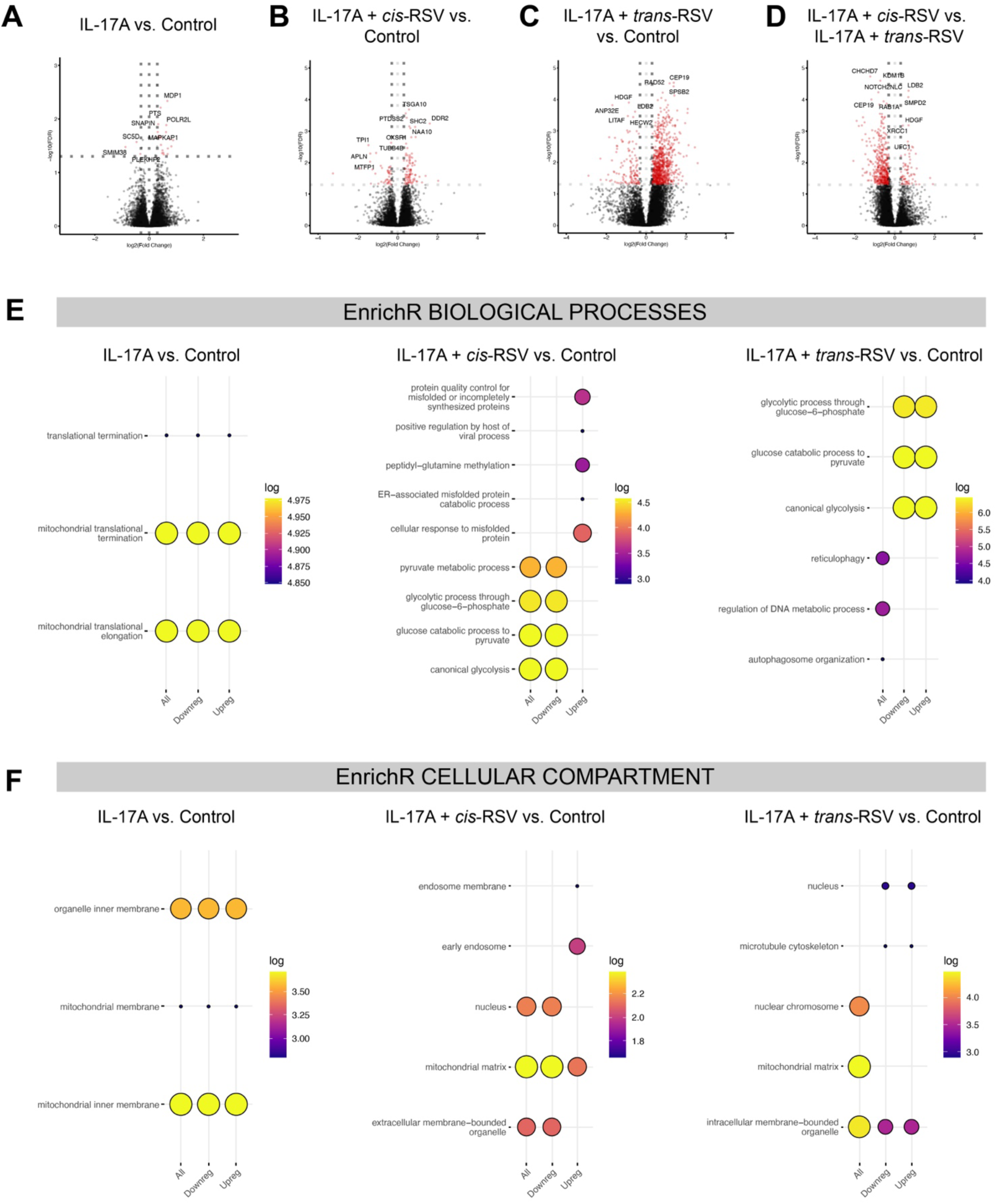
IL-17A, *cis-* and *trans*-RSV alter transcriptional programs associated with translation, mitochondrial function, and metabolism. (**A-D**) Volcano plots showing DEGs in human neurons treated with either IL-17A (**A**), IL-17A + *cis-*RSV (**B**) and IL-17A +*trans-*RSV (**C**), compared to control or comparing IL-17A + *cis*-RSV vs. IL-17A + *trans-*RSV (**D**). Log_2_ fold changes (LFC) gene expression (x-axis) and −log_10_ adjusted P values (y-axis) generated from DESeq2 differential gene expression analysis are shown. Vertical dotted lines represent 0.58 LFC (1.5 FC) and horizontal dotted line shows adjusted P=0.05. Significant DEGs are shown in red with the top DEGs labelled in the plot. (**E-F**) Functional enrichment analysis by EnrichR for biological process (**E**), and cellular compartment (**F**), show enrichment for all DEGs, upregulated and downregulated DEGs in IL-17A vs Control neurons or IL-17A + *cis-*RSV vs. control neurons or IL-17A + *trans-*RSV vs. control neurons. Circle size represents the number of DEGs in that category and the color represents the adjusted P value.

Functional enrichment analysis showed an overrepresentation of genes involved in translation and mitochondrial function in neurons treated with IL-17A alone. In contrast, neurons treated with either IL-17A + *cis-*RSV or IL-17A + *trans-*RSV showed enrichment in metabolic processes, particularly glycolysis and pyruvate metabolism (**Fig. 4E, Supplementary Fig. S3, and Supplementary Table S2**). Additionally, pathways associated with the endoplasmic reticulum (ER) and autophagosome function were enriched in neurons treated with IL-17A + *trans*-RSV (**Fig. 4F, Supplementary Fig. S3, and Supplementary Table S3**). Disruption of mitochondrial function, translation, or autophagy have been previously implicated in the pathobiology of ASD, suggesting the mechanistic relevance of our findings in relation to inflammatory pathways affecting neurodevelopment.

### WGCNA reveals additional regulatory nodes associated with the effect of inflammation on human neurons

To gain further insights into the transcriptional influence of IL-17A, *cis-*RSV, and *trans-*RSV at a system level, we performed weighted gene co-expression network analysis (WGCNA) (**Fig. 5, Supplementary Fig. S4, and Supplementary Table S4**). WGCNA identified several eigengene networks associated with each treatment, distinct from each other and from the control (**Fig. 5A and Supplementary Fig. S4**). We also investigated the correlation between eigengene and single-cell expression and disease association (**Supplementary Fig. S5**). The Allen Brain Atlas single-cell datasets (Tasic *et al*, 2018) showed enrichment of genes expressed in excitatory and inhibitory neurons (green, light cyan, midnight blue and yellow modules), as well as in oligodendrocyte precursors (OPCs) and astrocytes (blue module) (**Fig. 5B**). To examine the potential relationship between dysregulation in neuropsychiatric disorders and alterations in transcriptional profiles driven by inflammation, we used published meta-analysis of neuropsychiatric brain gene expression from the PsychENCODE Consortium (Wang *et al*, 2018). We found that there are modules associated with ASD (blue, pink, and yellow modules), schizophrenia (blue and pink modules), and bipolar disorder (black, pink, red, and yellow modules) (**Fig. 5C**). Next, we conducted functional gene enrichment analysis on the WGCNA datasets to determine whether specific eigengene networks associated with disease phenotypes were enriched in specific functional categories (**Fig. 5D, Supplementary Fig. S4 and Supplementary Table S3-S4**). Functional gene enrichment analysis of the WGCNA datasets revealed significant enrichment in eigengene networks associated with translation (royal blue module), autophagy (turquoise module), and RNA processing (blue module). These findings highlight the relevance of inflammatory pathways in modulating gene networks during early brain development, emphasizing their potential impact on neuropsychiatric disease mechanisms.

**Figure 5.**
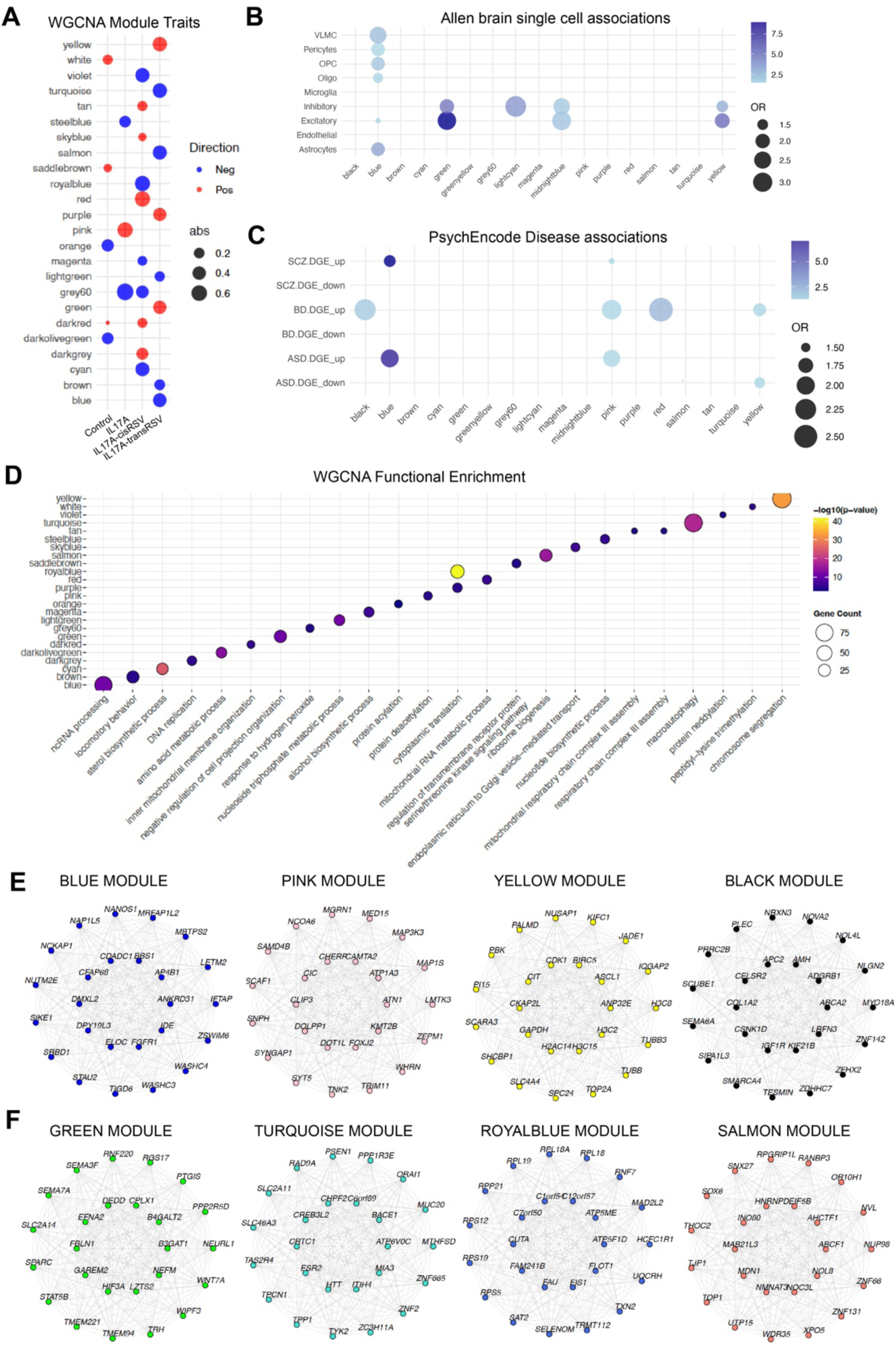
Disease association and functional enrichment by WGCNA analysis. (**A**) WGCNA module trait relationships are shown for control, IL-17A, IL-17A + *cis-*RSV and IL-17A + *trans-*RSV samples. Each module is represented with the direction of gene expression change, indicated as either positive (red) or negative (blue). (**B**) Magma plots show WGCNA enrichment for single cell expression from the Allen brain atlas. (**C**) WGCNA modules showing disease association with neuropsychiatric disorders highlights upregulated and downregulated DEGs within each module. The size of the solid black circles corresponds to the number of DEGs, and the color intensity of blue represents the log P value, with darker hues indicating greater significance. (**D**) WGCNA pathway enrichment analysis is shown across the different modules with colors depicting the log P value, indicating the significance of the enriched pathways. (**E-F**) Gene networks are shown with relevant hub genes for the blue, pink, yellow, and black modules (**E**), and for the green, turquoise, royalblue, and salmon modules (**F**).

### Impact of inflammation on gene expression in ASD: linking translation and immune response

Based on the transcriptional effect of IL-17A as well as *trans*-RSV on human cortical excitatory neurons we observed, we decided to examine the extent to which these transcriptional changes might overlap with molecular signatures associated with ASD patients. Therefore, we selected a group of ASD patients that presented with a larger head circumference and in some instances were exposed to infections postnatally or in utero, hereafter referred to as ASDi (**Supplementary Table S5**). To begin to understand the connection between inflammation and ASD pathogenesis in unaffected and ASD subjects, peripheral blood mononuclear cells (PBMCs) were collected and transformed into lymphoblastoid cells and subjected to metabolic profiling and transcriptome analysis. Metabolic profiling was conducted on lymphoblastoid cells using different cytokines (IL-6, IL-1β, IL-8, TNF-α and IFN-γ) that have been previously associated with ASD (**Fig.6A**). Metabolic profiling revealed decreased responsiveness to certain cytokines (IL-6, IL-1β, IL-8, IFN-γ) in ASD lymphoblasts compared to controls, with no significant difference for TNF-α. To explore the potential mechanisms that could underlie the response to the different cytokines we conducted a transcriptome analysis of a subset of the ASDi cohort that in addition to sensitivity to disease also had a larger head circumference (**Supplementary Table S5**). Transcriptome analysis in ASDi lymphoblastoid cells identified distinct gene expression signatures compared to controls, with upregulated DEGs involved in chromosome segregation and cell division and downregulated DEGs related to translation and immune function (**Fig. 6B-C, Supplementary Fig. S5, Supplementary Table S6-S7**). Gene ontology analysis highlighted enrichment for pathways controlling translation, cell division, as well as endoplasmic reticulum function (**Fig. 6C and Supplementary Table S7**). Additionally, functional enrichment analysis revealed upregulated RNA binding activity among the DEGs, suggesting a significant role of RNA processing in the response to inflammation (**Fig. 6D**). Similarly, molecular function analysis by EnrichR revealed a major node comprised by upregulated DEGs associated with RNA binding (**Supplementary Fig. S5 and Supplementary Table S7**). Additionally, KEGG pathway analysis identified several viral response pathways, in addition to splicing and translation signatures (**Fig. 6E**). Finally, disease analysis showed enrichment for Schizophrenia and Parkinson’s disease among the upregulated DEGs (**Fig. 6F and Supplementary Table S6**). Therefore, our transcriptomic studies in human neurons and in ASDi patient lymphoblastoid cells suggest that dysregulation of translation might be a common molecular signature associated in some instances with ASD cases that present with immune dysfunction and potentially with the effect of MIA in the developing brain.

**Figure 6.**
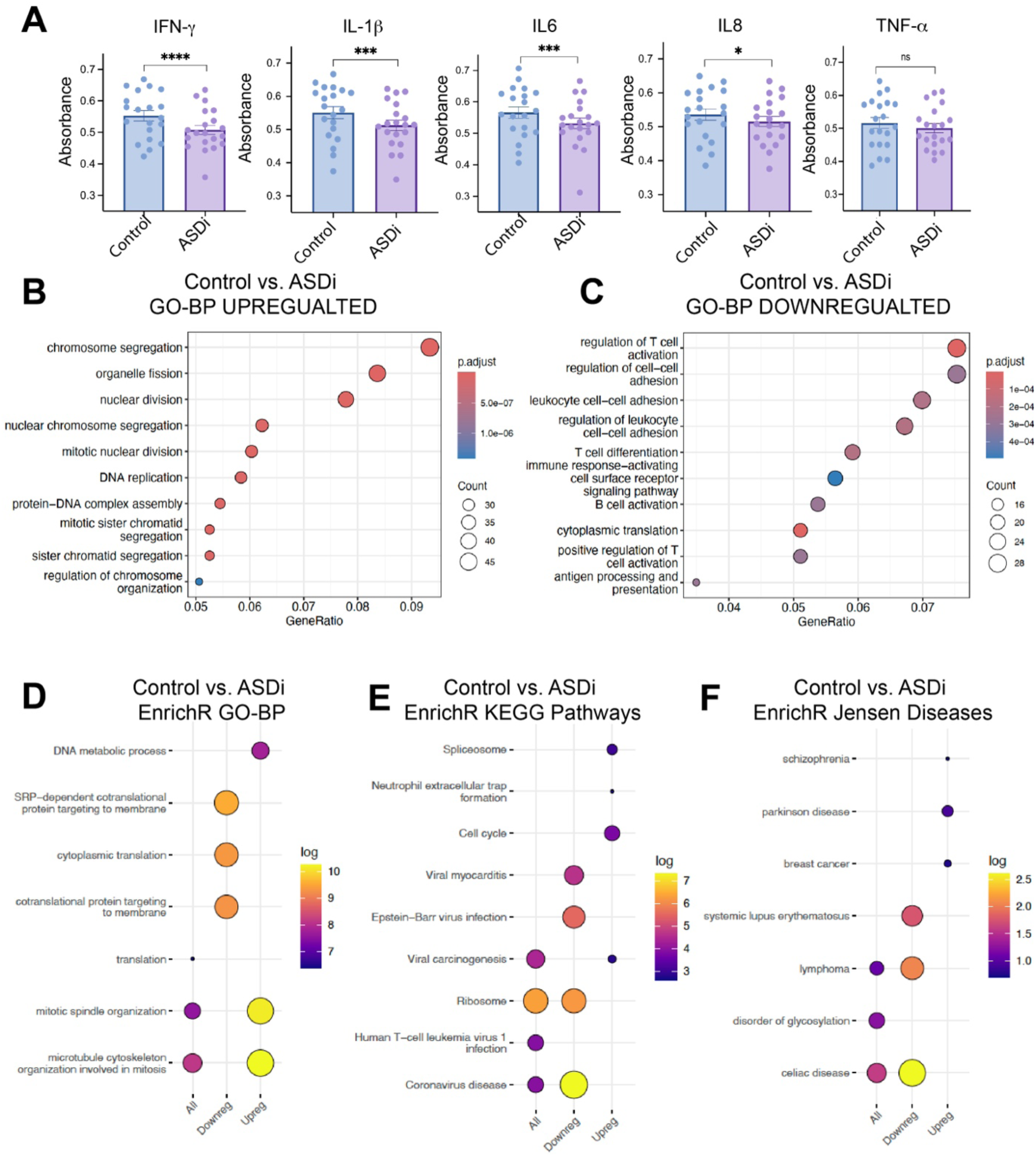
Metabolic and transcriptome analysis of ASD patient lymphocytes. (**A**) Graphs show biolog measurements for control (blue bars and circles) and ASDimmune (ASDi) (purple bars and circles) patient lymphocytes treated with different cytokines. Mean + SEM are shown with individual circles representing the average across multiple measurements. Data was analyzed as grouped data using two-way ANOVA. *P <0.04, ***P < 0.0004, and ****P < 10e^-15^. (**B-C**) Gene enrichment analysis using gene ontology (GO) for biological process (BP) for upregulated (**B**) and downregulated DEGs (**C**). (**D-F**) Functional enrichment analysis by EnrichR for biological process (**D**), KEGG pathways (**E**), and Jensen diseases (**F**) show enrichment for all DEGs, upregulated and downregulated DEGs in Control vs. ASDi lymphocytes. Circle size represents the number of DEGs in that category and the color represents the adjusted P value.

## DISCUSSION

The effects of MIA on the fetal developing brain are increasingly documented across multiple animal models (Vacharasin *et al*, 2024). However, mechanistic understanding of human-specific mechanisms underlying the impact of inflammation on the fetal brain remains limited. The advent of stem cell technology in combination with two- and three-dimensional neural models is now allowing us to dissect the complex immune-neuro axis in order to understand how increased inflammation could affect human fetal brain development (Freitas *et al*, 2018). Our studies using hESC-derived cortical excitatory neurons uncover a novel mechanism linking inflammation to increased DNA damage and the subsequent activation of the cGAS-STING pathway. While the cGAS-STING dysfunction has been previously implicated in neuroimmune and neurodegenerative diseases (Li *et al*, 2024), its role in ASD pathogenesis has not been previously reported. Increasing evidence implicates proteins involved in the DNA damage response in multiple neurodevelopmental disorders (Cunningham *et al*, 2024). However, the connection between increased DNA damage with cGAS-STING activation in response to inflammatory signals and autism pathogenesis in human neurons has not been previously reported. The connection between DNA damage, cGAS-STING activation, and autism pathogenesis in human neurons underscores a critical gap in the current understanding, highlighting the need for more specific human models to study neurodevelopmental disorders. More importantly, our work further suggests that the increased DNA damage could be in part associated with increased oxidative stress, as indicated by increased nuclear translocation of TyrRS in human neurons treated with inflammatory signals. Under oxidative stress, nuclear translocation of TyrRS will suppress protein synthesis and protect against DNA damage (Cao *et al*., 2017; Jones *et al*, 2023; Wei *et al*., 2014). This mechanism is crucial because it points to a potential pathway through which oxidative stress could exacerbate DNA damage in neurons, linking it to neurodevelopmental disorders like ASD. The dysregulation of gene programs controlling translation in response to inflammation, observed in both human neurons and ASD patient lymphoblasts, suggests a broader impact on cellular function in response to inflammatory signals, potentially affecting the neuro-immune axis in ASD.

Our work also highlights the importance of mitochondrial function during brain development, as a major transcriptomic signature we observed was altered in human neurons treated with different inflammatory cues. Modulation of Ca+ signaling and ATP production by mitochondria are essential for synapse development and plasticity (Khaliulin *et al*, 2024) and could be underlying the defects in spontaneous neuronal activity we observed in neurons treated with IL-17A and *trans-*RSV. These disruptions may contribute to the defects in spontaneous neuronal activity observed in treated neurons, suggesting that mitochondria play a crucial role in modulating the response to inflammation during development. Additionally, we identified robust metabolic changes in response to inflammatory signals, particularly in glucose metabolism, indicating a metabolic reprogramming akin to what is seen in immune cells (Mouton *et al*, 2023). This suggests that neurons respond to inflammation by altering their metabolic processes, which could influence cellular functions critical for brain development.

In conclusion, our study provides new insights into how inflammation disrupts molecular pathways in human neurons, particularly highlighting the connection between DNA damage, oxidative stress, and immune response. The involvement of the cGAS-STING pathway and mitochondrial dysfunction in response to inflammatory signals offers a deeper understanding of ASD pathogenesis and points to potential therapeutic targets for intervention. By distinguishing the effects of different resveratrol isomers—*cis*-RSV as potentially protective against inflammation—this work emphasizes the need for targeted therapies that can modulate these pathways in developing neurons. Future research will continue to explore these mechanisms in greater detail, particularly as they relate to the development of therapeutic strategies for neurodevelopmental disorders affected by inflammation.

## MATERIALS AND METHODS

### Stem cell culture

Human embryonic stem cells (ESC) H9 were obtained from WiCell and were cultured as previously described (Lizarraga *et al*, 2021). ESC colonies were inspected daily to ensure there was no spontaneous differentiation. ESCs were maintained in mTeSR™ Plus basal medium supplemented with 5X supplement (STEMCELL Technologies, #100-0276) seeded on Matrigel (Corning #354277) coated plates. ESCs were fed daily and were passaged every 3-4 days at a 1:6 ratio using ReLeSR™ (STEMCELL Technologies, #100-0483) following manufacturer protocol. To minimize chromosomal abnormalities, ESCs were maintained only up to passage 50.

### Generation of forebrain cortical excitatory neurons from human ESCs

H9 ESCs were differentiated into forebrain cortical excitatory neurons as described previously (Lizarraga *et al*., 2021; Shi *et al*., 2012). Briefly, ESC colonies were dissociated with Gentle Cell Dissociation Reagent (STEMCELL Technologies, #100-0485) and were plated in mTeSR™ Plus medium (STEMCELL Technologies, #100-0276) supplemented with ROCK inhibitor Y27632 (STEMCELL Technologies, #72304) at 1x 10^6^ cells per well into 12-well tissue culture plates coated with Matrigel (Corning #354277). After twenty four hours, a complete monolayer should have formed and cell media was replaced with neural induction medium (NIM) made with 1:1 DMEM/F-12 GlutaMAX (Thermo Fisher Scientific, #10565018) and Neurobasal media (Thermo Fisher Scientific, #21103049) supplemented with: N2 (Thermo Fisher Scientific #17502-048), B27 (Thermo Fisher Scientific, #17504044), 5µg/ml insulin (Sigma-Aldrich, #I0526-5ML), 1mM L-glutamine (Thermo Fisher Scientific, #25-030-081), 100 μM nonessential amino acids (Thermo Fisher Scientific, #11-140-050), 100µM β-mercaptoethanol (Sigma-Aldrich, #M6250-100ML), 50U/ml penicillin-streptomycin (Thermo Fisher Scientific, #15-140-122), 1µM Dorsomorphin (STEMCELL Technologies,#72102) and 10µM SB431542 (STEMCELL Technologies, #72234). Neuronal progenitors were expanded in NIM with the addition of 10µM recombinant FGF2 (PeproTech, #100-18C) from days 17 to 21. After day 21, neurons were fed with half media changes every other day using neural maintenance medium (NMM), containing all the components of NIM except for Dorsomorphin and SB431542. For subsequent experiments neurons were seeded on either 8-well chamber slides or multi-well tissue culture plates coated with 20µg/ml laminin (Sigma-Aldrich, #L2020-1MG) and poly-ornithine (Sigma-Aldrich, #P4957-50ML).

### Western blotting

For western blot analysis, neurons at day 55 of neuronal induction were washed twice with cold 1X PBS and lysed in cell lysis buffer (20 mM Tris-HCl (pH 7.5), 150 mM NaCl, 1 mM Na_2_EDTA, 1 mM EGTA, 1% Triton, 2.5 mM sodium pyrophosphate, 1 mM beta-glycerophosphate, 1 mM Na_3_VO_4_, 1 µg/ml leupeptin supplemented with protease inhibitor). The lysates were centrifuged at 12,000 *g* for 15 min at 4 °C. Lysates were quantified using Pierce BCA Protein Assay kit (Thermo Scientific #23227), and equal amounts of protein were loaded onto a 4 to 12% gradient gel (NuPAGE Invitrogen). Protein was transferred from the gel to 0.2 μm NC membranes at 25 V for 10 min using transfer stacks (iBlot3 Invitrogen) and blocked with LiCor TBS blocking buffer (#927-60003) for 1 hour at room temperature before application of primary antibodies. Primary and secondary antibodies were incubated overnight at 4 °C and for 1 hour at room temperature, respectively. Membranes were incubated with primary antibodies using the following dilutions: cGAS (Proteintech, #26416-1-AP 1:500), STAT3 (Cell Signaling Technology, #9139S, 1:500), phospho-STAT3 (Cell Signaling Technology, #9145S,1:500), NF-κB (Cell Signaling Technology, #8242S, 1:500), NF-κB2 (Cell Signaling Technology, #3017S, 1:500), TyrRS (Abcam, #ab50961, 1:1000), TTL (Proteintech, #66076-1-IG, 1:1000), Tyrosinated tubulin (Millipore Sigma, #MAB1864-I,1:2000), MAPK (Cell Signaling Technology, #9102S, 1:1000), p-MAPK (Cell Signaling Technology, #9101S, 1:1000), and GAPDH (Cell Signaling Technology, #97166S, 1:1000) prepared in LiCor Intercept T20 (TBS) Antibody Diluent (#927-65003). Blots were washed 1X TBST (10 mM Tris-HCl pH 8.0, 150 mM NaCl, 0.01% Tween-20) after primary antibody incubation. Fluorescent secondary antibodies diluted at 1:5000, were used for anti-rabbit (Thermo Scientific, #A32729), anti-mouse (Thermo Scientific, #A32735), and anti-rat (Thermo Scientific, #A11006). Blots imaged using LiCor Odyssey CLX scanner were quantified using ImageJ (Version 1.53c).

### Pharmacological Treatments

Stock solutions for RSV were prepared at a concentration of 1000x in ethanol and then diluted into cell culture media to achieve the desired final concentration. *cis-*RSV was purchased from Cayman Chemicals (#10004235, ≥ 98% purity), dissolved in ethanol to prepare a stock solution at 100 mM. *trans-*RSV was obtained from Millipore-Sigma (#34092, ≥ 99% purity) and also dissolved in ethanol to make a 100 mM stock solution. IL-17A was purchased as a lyophilized powder from R&D Systems (#317-ILB-050), dissolved in PBS to create a stock solution of 50 µg/ml, and stored at −80°C until required for use. For all experiments, the treatments were done using 50 µM for RSV isomers and 50ng/ml for IL-17A.

### Immunocytochemistry

At day 32 of neuronal induction, ESC-derived human cortical neurons were plated (30,000 cells/well) in 8-well chamber slides (CELLTREAT, #229168) coated with 0.01% poly-L-ornithine and 20µg/ml laminin. For each experiment, three independent neuronal inductions were used. At day 55, neurons were treated with either DMSO, IL-17A, *cis*-Resveratrol (*cis-*RSV), *trans*-Resveratrol (*trans*-RSV), IL-17A + *cis-*RSV or IL-17A + *trans-*RSV and after 4 hours neurons were briefly rinsed with 1XPBS (Thermo Scientific, #70011044), fixed with 4% paraformaldehyde (Electron Microscopy Sciences, #15710-S) for 15 minutes (min) at room temperature, washed three times for 5 min each with 1XPBS, permeabilized with 0.25% Triton X-100 (Sigma-Aldrich, #X100-100ML) for 10 min at room temperature, washed three times for 5 min each with room temperature 1X PBS, and blocked with 10% goat serum solution (Thermo Scientific, #50062Z) for one hour at room temperature. After blocking, primary antibodies against either MAP2 (Abcam #36592 1:1250), NF-κB (Cell Signaling Technology, #8242S, 1:400), cGAS (Proteintech, #26416-1-AP, 1:200), Tyrosinated Tubulin (Millipore Sigma, #MAB1864-I, 1:500), TyrRS (Abcam, #ab50961, 1:50) diluted in 2% goat serum and 0.2% Triton X-100 (Lizarraga *et al*., 2021) were applied to cells and incubated overnight at 4°C. Following primary antibody treatment, cells were washed three times with 1XPBS for 5 minutes each time, and then incubated for 1 hour at room temperature with secondary antibodies diluted in 1XPBS with 2% goat serum (AlexaFluor-594 anti-mouse Thermo Fisher Scientific #A11032, Alexa Fluor-647 anti-rabbit Thermo Fisher Scientific # A-21245, and Alexa Fluor-488 anti-chicken Thermo Fisher Scientific # A32931, all diluted at 1:1000). Cells were washed three times with 1XPBS, and slides were mounted with ProLong Glass Antifade Mountant with DAPI (Thermo Scientific #P36985) under 1 mm rectangular coverslips (Fisher Scientific, #12-548-5P).

### Image acquisition and analysis

#### Imaging

Images were captured using an FV3000 Olympus Confocal Microscope equipped with a 60X or a 20X objective lens. Fluorescent images of 30 to 35 random cells were taken for each experimental condition for at least three independent experiments.

#### Protein levels intensity analysis

Protein colocalization was assessed for two different fluorophores by generating a maximum intensity z-projection from image stacks. Regions of interest (ROIs) were created around individual cells using the select tool in Fiji ImageJ, and individual masks were generated to exclude external signals. The ROI was duplicated in split color channels—red (NF-κB, TyrRS, cGAS) and blue (DAPI) or cyan (MAP2) across different images. Pixel intensity values were thresholded across all images to ensure consistency in the analyzed areas and to exclude background noise. The software calculated the mean gray value, representing the average pixel intensity within the selected area. These measurements were grouped per independent neuronal induction and were compared for analysis.

#### γ-H2A.X foci analysis

Number of foci per cell were measured for each condition with 30 to 35 nuclei analyzed per condition for each experiment. First, the regions of interest (ROIs) around individual cell nuclei were defined using DAPI staining to mark nucleus boundaries. The analyze Particles function in ImageJ was used to measure γ-H2A.X distribution. After thresholding the images to a uniform value across samples, the number of particles per ROI were counted, representing the number of foci per cell. These measurements were grouped per independent neuronal induction and were compared for analysis.

### Library preparation and sequencing

RNA was extracted using the RNeasy Micro Kit (Qiagen, #74004) following the manufacturer’s instructions. For each sample, RNA quality was analyzed using Agilent 2200 Tape Station system. cDNA libraries were prepared with the Collibri™ Stranded RNA Library Prep Kit for Illumina™ Systems (Invitrogen, #A38996096). Sequencing was performed using 150-bp reads using paired-end chemistry on an Illumina NovaSeq, to achieve a depth of 100 million reads.

### RNA-seq data processing and analysis

We assessed the quality of raw sequence data using FastQC (version 0.11.9) and TrimGalore (version 0.6.6). Raw reads were mapped to the Ensembl reference genome 104 (GRCh38) using the transcript-level quantifier Salmon (version 1.5.2) in mapping-based mode. Count matrices were generated from transcript-level quantification files using the tximport package (version 1.30.2). All RNAseq data is publicly available through GEO #. Differentially expressed genes (DEGs) were identified with DESeq2 (version 1.32.0) (Love *et al*, 2014), considering genes with an adjusted p-value < 0.05 and a fold change ≥ |1.5| as statistically significant. Pathway enrichment analysis (GO, KEGG, DisGeNET) was performed using the EnrichR (version 3.2) and clusterProfiler (version 4.10.1) package in R.

### Co-expression network analysis

To identify modules of co-expressed genes in the RNA-seq data, we carried out weighted gene co-expression network analysis (WGCNA v1.69)(Langfelder & Horvath, 2008). A soft-threshold power was automatically calculated to achieve approximate scale-free topology (R^2^>0.85). Networks were constructed with *blockwiseConsensusModules* function with biweight midcorrelation (bicor). The modules were then determined using the dynamic tree-cutting algorithm. To ensure robustness of the observed network, we used a permutation approach recalculating the networks 200 times and comparing the observed connectivity per gene with the randomized one. None of the randomized networks showed similar connectivity, providing robustness to the network inference. Module sizes were chosen to detect small modules driven by potential noise on the adjusted data. Deep split of 4 was used to more aggressively split the data and create more specific modules. Spearman’s rank correlation was used to compute module eigengene – memory oscillatory signature associations.

#### Functional Enrichment

The functional annotation of the genes within the modules was performed using GOstats (v2.56.0) using the GO and KEGG databases. Expressed genes (13818) were used as background. A one-sided hypergeometric test was performed to test overrepresentation of functional categories and Benjamini-Hochberg adjusted p-value used for multiple comparisons correction.

#### Neuropsychiatric genes

Autism spectrum disorders associated genes used for **Supplementary Fig. S5E** were downloaded from SFARI database (Banerjee-Basu & Packer, 2010). Modules and genes differentially expressed in ASD, SCZ, and BD were downloaded from an independent source(Gandal *et al*, 2018). Differentially expressed cell-type markers from single-nuclei RNA-seq of ASD were downloaded from published dataset (Velmeshev *et al*, 2019).

#### GWAS data and enrichment

We used genome-wide gene-based association analysis implementing MAGMA (v1.07) (de Leeuw *et al*, 2015). We used the 13818 protein-coding genes from human gencode v38 as background for the gene-based association analysis. Suplementary table # reports MAGMA statistics for each of the GWAS data analyzed. GWAS acronyms were used for the figures (e.g.*, ASD = Autism Spectrum Disorders, BD = Bipolar disorder, SZ = Schizophrenia).* Gene set enrichment was applied to correlated genes. We used a Fisher’s exact test in R with the following parameters: alternative = “greater”, conf.level = 0.95. We reported Odds Ratios (OR) and Benjamini-Hochberg adjusted P-values (FDR).

*Transcriptomic data available upon request*.

### Metabolic profiling (PM-M) of Lymphoblastoid cell lines

A specialized technology called Phenotype Mammalian MicroArray (PM-M) plates was developed by Biolog (Hayward, CA, USA) to identify metabolic signatures of behavioral symptoms in PMS. To obtain a metabolic profile, lymphoblastoid cell lines (LCLs) were established from peripheral blood samples collected by venipuncture, using lymphocyte immortalization via Epstein–Barr virus (Srikanth *et al*, 2021). LCLs were harvested in Sigma RPMI-1640 with 15% fetal bovine serum (FBS) from Atlanta Biological (Flowery Branch, GA, USA), two mM L-Glutamine, 100 U/mL Penicillin, and 100 μg/mL Streptomycin from Sigma-Aldrich (St. Louis, MO, USA).

The PM-M plates measure the cellular production of NADH (nicotinamide adenine dinucleotide, reduced form) in the presence of different compounds to assess metabolic activity. Microplates are utilized with diverse molecules, acting as energy sources (plates PM-M1 to M4) or as metabolic effectors (plates PM-M5 to M8). A single compound is within each well, and a colorimetric redox dye chemistry monitors the production of NADH per well. The energy sources include carbohydrates, carboxylic acids, ketone bodies, and nucleotides in plate PM-M1, as well as amino acids (both alone and as dipeptides) in plates PM-M2 to M4. The metabolic effectors include ions in PM-M5, along with cytokines, growth factors, and hormones in PM-M6 to M8. PM-M plates were incubated with 20,000 lymphoblastoid cells per well in a volume of 50 μL, using the modified Biolog IF-M1 medium. Media for plates PM-M1 to M4 were prepared by adding the following to 100 mL of Biolog IF-M1: 1.1 mL of 100× penicillin/streptomycin solution, 0.16 mL of 200 mM Glutamine (final concentration 0.3 mM), and 5.3 mL of fetal bovine serum (final concentration 5%). Cells were incubated for 48 h at 37°C in 5% CO2. After this first incubation, Biolog Redox Dye Mix MB was added (10 μL/well) and plates were incubated under the same conditions for an additional 24 h, during which time cells metabolize the sole carbon source in the well. As the cells metabolize the carbon source, tetrazolium dye in the media is reduced, producing a purple color according to the amount of NADH generated. During the 24 h of exposure to dye, plates were incubated in the Omnilog system, which measured the optical density of each well every 15 minutes, generating 96 data points. At the end of the 24-h incubation, plates were analyzed utilizing a microplate reader with readings at 590 and 750 nm. The first value (A_590_) indicated the highest absorbance peak of the redox dye and the second value (A_750_) gave a measure of the background noise. The relative absorbance (A_590-750_) was calculated per well as previously described (Boccuto *et al*, 2013). The relative absorbance for 20 control and ASD cell lines after treatment with different cytokines was analyzed using one-way ANOVA analysis in Graphpad prism software.

### Multielectrode Array Analysis

Neurons were seeded on Cytoview48 MEA plates (Axion Biosystems, #M768-tMEA-48W) at a density of 100,000 cells per well on day 40-41 of neuronal differentiation. Plates were pre-coated using poly-D-lysine (100μg/ml) and laminin (20μg/ml) for 1 hour each at 37°C following the standard coating protocol. Neurons were plated in neural maintenance medium (NMM) supplemented with laminin (20μg/mL). After 24 hours, the medium was replaced with BrainPhys medium (STEMCELL Technologies, **#**05790) containing NeuroCult™ SM1 and N2 supplements (STEMCELL Technologies, **#**05711 and #07152), dibutyryl cyclic-AMP (500μg/ml; STEMCELL Technologies, # 100-0244), GDNF (Peprotech, #450-10) and BDNF (Peprotech, 450-02) both 20 ng/ml, and ascorbic acid (200nM, STEMCELL Technologies, # 72132). Neurons were maintained on MEA plates with half-medium changes every third day until the experiment. On day 55 of differentiation, treatments were applied to eight wells per experimental condition. Following treatment, electrophysiological activity was recorded using the MEA Maestro Pro system (Axion Biosystems) under standard neural activity conditions. Recordings were taken every 2 hours for 15 minutes over a 24-hour period. Throughout the experiment, cells were maintained at 37°C in 5% CO₂. Spike counts and mean-weighted firing rates were obtained from compiled statistics files for each recording. The 24-hour experimental results were analyzed collectively for each condition, grouping data across all time points to assess overall trends and differences.

## ACKNOWLEDGEMENTS

The authors wish to thank: Current members of the Lizarraga laboratory for critical reading of the manuscript and previous members of the Lizarraga laboratory, Dr. Pankaj Ghate and Dr. Janay Vacharasin for their preliminary observations; the families that participated in the clinical studies at Greenwood Genetics Center (GCC); Dr. Roger Stevens and Dr. Richard Steet at GGC for the generous donation of control and patient lymphoblastoid cells for functional studies in the Lizarraga laboratory; Cyndi Skinner and Jessica Coleman at GGC who handled the patient samples and clinical histories; and Dr. Stefano Berto (Medical University of South Carolina) for sharing WGCNA bioinformatics pipelines. The research reported in this publication was supported by the National Institute of Mental Health of the National Institutes of Health under Award Number R01MH127081 to SBL. The content is solely the responsibility of the authors and does not necessarily represent the official views of the National Institutes of Health.

## AUTHORS CONTRIBUTIONS

SBL and MJ conceived the study, oversaw the experimental design and data analysis, and wrote the manuscript. SBL led the funding acquisition. MJ conducted the majority of the experiments and data analysis. CK and AG contributed to the biochemical and imaging analysis. LB conducted the metabolic experiments. BY contributed to the RNAseq analysis. SM provided advice on DNA damage studies. All authors contributed to editing the manuscript.

