## Supplemental Material for "The intersection of inflammation and DNA damage as a novel axis underlying the pathogenesis of autism spectrum disorders"

### SUPPLEMENTARY MATERIAL

Megha Jhanji [1,2], Colleen L. Krall [1,2], Alexis Guevara [1,2], Brian Yoon [1,2], Mathew Sajish [3], Luigi Boccuto [4,5], and Sofia B. Lizarraga [1,2&]

[1] Department of Molecular Biology, Cell Biology and Biochemistry, Brown University, Providence, RI
[2] Center for Translational Neuroscience, Carney Brain Institute, Brown University, Providence, RI
[3] Department of Drug Discovery and Biomedical Sciences, College of Pharmacy, University of South Carolina, Columbia, SC
[4] JC Self Research Institute, Greenwood Genetic Center, Greenwood, SC
[5] School of Nursing, College of Behavioral, Social, and Health Sciences, Clemson University, Clemson, SC

*Running title:* *Inflammatory pathways induce DNA damage response in human neurons*

### SUPPLEMENTARY FIGURES

#### Supplementary Figure S1 related to main Figure 1

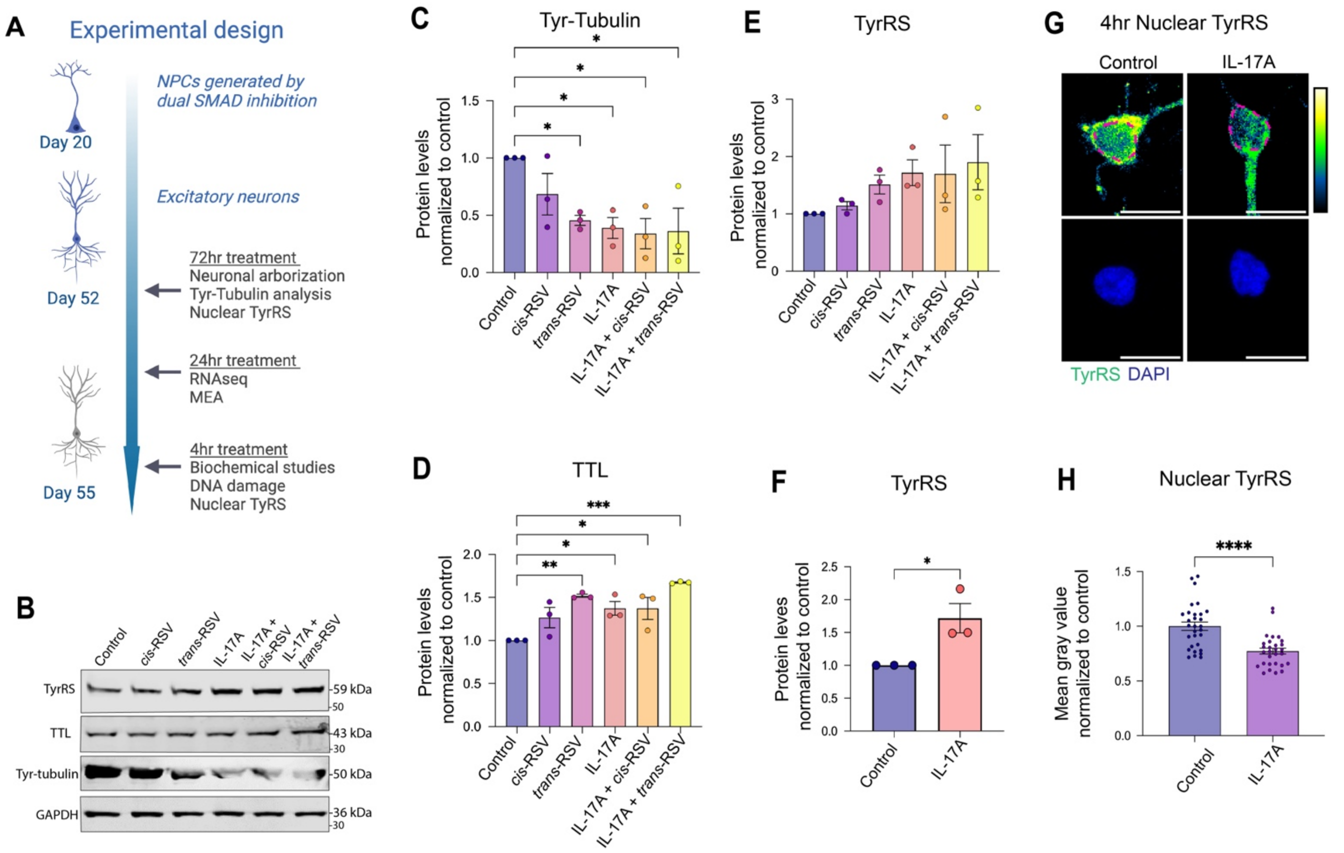


**Supplementary Figure S1. Experimental design and total protein analysis of human neurons.** (**A**) Experimental design is shown for all experiments with human neurons. (**B**) Representative image of western blot utilized to analyze changes in protein levels in response to different treatments. (**C-E**) Graphs depict protein quantification normalized to loading control and to untreated condition for at least 3 independent experiments. Mean ± SEM are shown with individual experiments represented by individual circles for control neurons (dark blue), neurons treated with cis-RSV (purple), trans-RSV (lilac), IL-17A (red), IL-17A+ cis-RSV (orange) and IL-17A + trans-RSV (yellow) for tyrosinated tubulin (tyr-Tubulin in C), tubulin tyrosine ligase (TTL in D), tyrosyl-tRNA synthetase (TyrRS in E). Statistical analysis was done using one way ANOVA with Dunnett's multiple comparisons test. *P < 0.05, ** P <0.001, and ***P <0.0001. (**F**) Re-analysis of TyrRS for control and IL-17A using student t test comparison, *P < 0.05. (**G**) Representative image of nuclear TyrRS after 4 hours of IL-17A treatment, with quantification shown as mean ± SEM and individual data points represented as circles for control (blue) and IL-17A (purple). Statistical analysis was performed using a two-tailed unpaired t-test (***P < 0.0001).

**Supplementary Figure S2 related to main Figure 3**

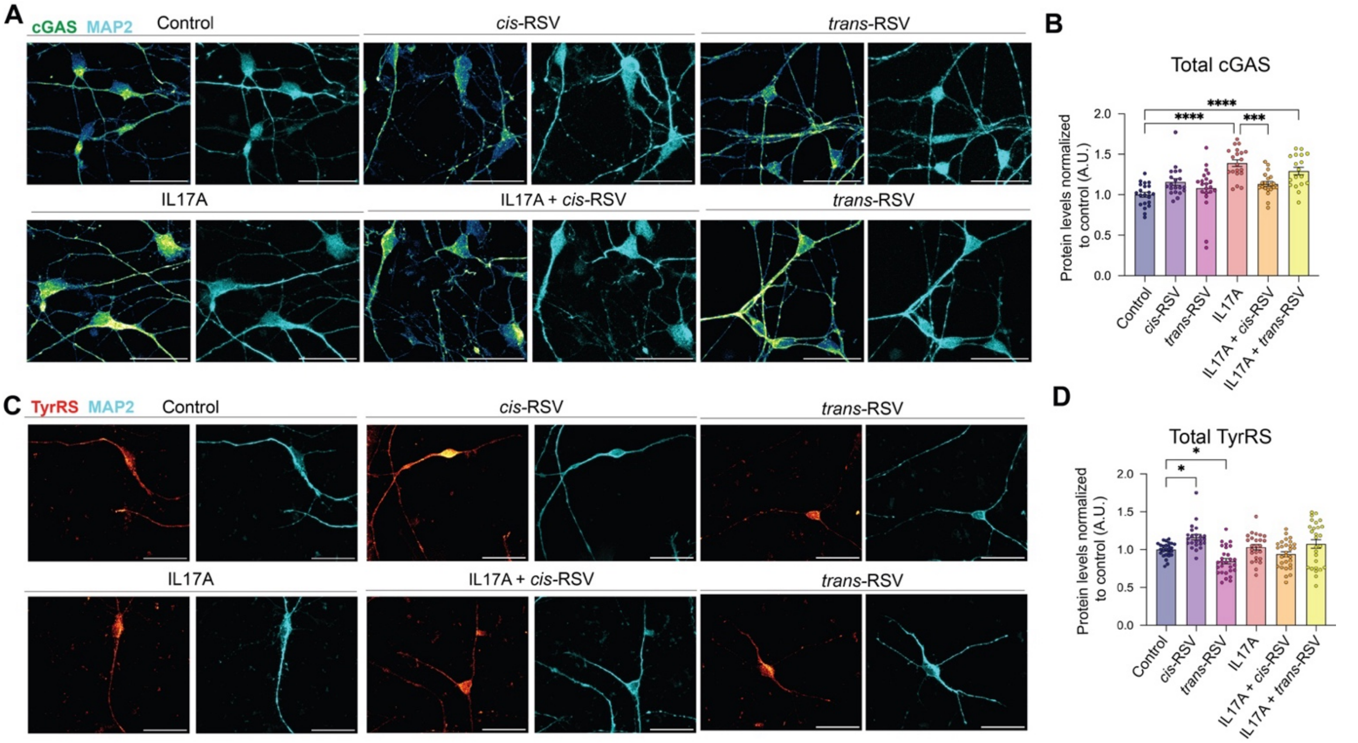


**Supplementary Figure S2. Analysis of total levels of cGAS and tyrRS in human neurons by confocal microscopy.** (**A**) Representative images are shown for the different treatments. Neurons were stained with antibodies against cGAS (green) and MAP2 (cyan). (**B**) Graphs depict mean gray value normalized to control neurons for at least 3 independent experiments and at least 30 neurons measured per experiment. Mean ± SEM are shown with individual circles representing the average across 3 experiments for control neurons (dark blue), neurons treated with *cis*-RSV (purple), *trans*-RSV (lilac), IL-17A (red), IL-17A+ *cis*-RSV (orange) and IL-17A + *trans*-RSV (yellow) for total cGAS. Statistical analysis was done using two way ANOVA with Sidak's multiple comparisons test, ***P <0.001, and ****P <0.0000001. (**C**) Representative images are shown for the different treatments. Neurons were stained with antibodies against TyrRS (red) and MAP2 (cyan). (**D**) Graphs depict mean gray value normalized to control neurons for at least 3 independent experiments and at least 30 neurons measured per experiment. Mean + SEM are shown with individual circles representing the average across 3 experiments for control neurons (dark blue), neurons treated with *cis*-RSV (purple), *trans*-RSV (lilac), IL-17A (red), IL-17A+ *cis*-RSV (orange) or IL-17A + *trans*-RSV (yellow) for total TyrRS. Statistical analysis was done using two-way ANOVA with Sidak's multiple comparisons test, *P <0.05.

**Supplementary Figure S3 related to main Figure 4**

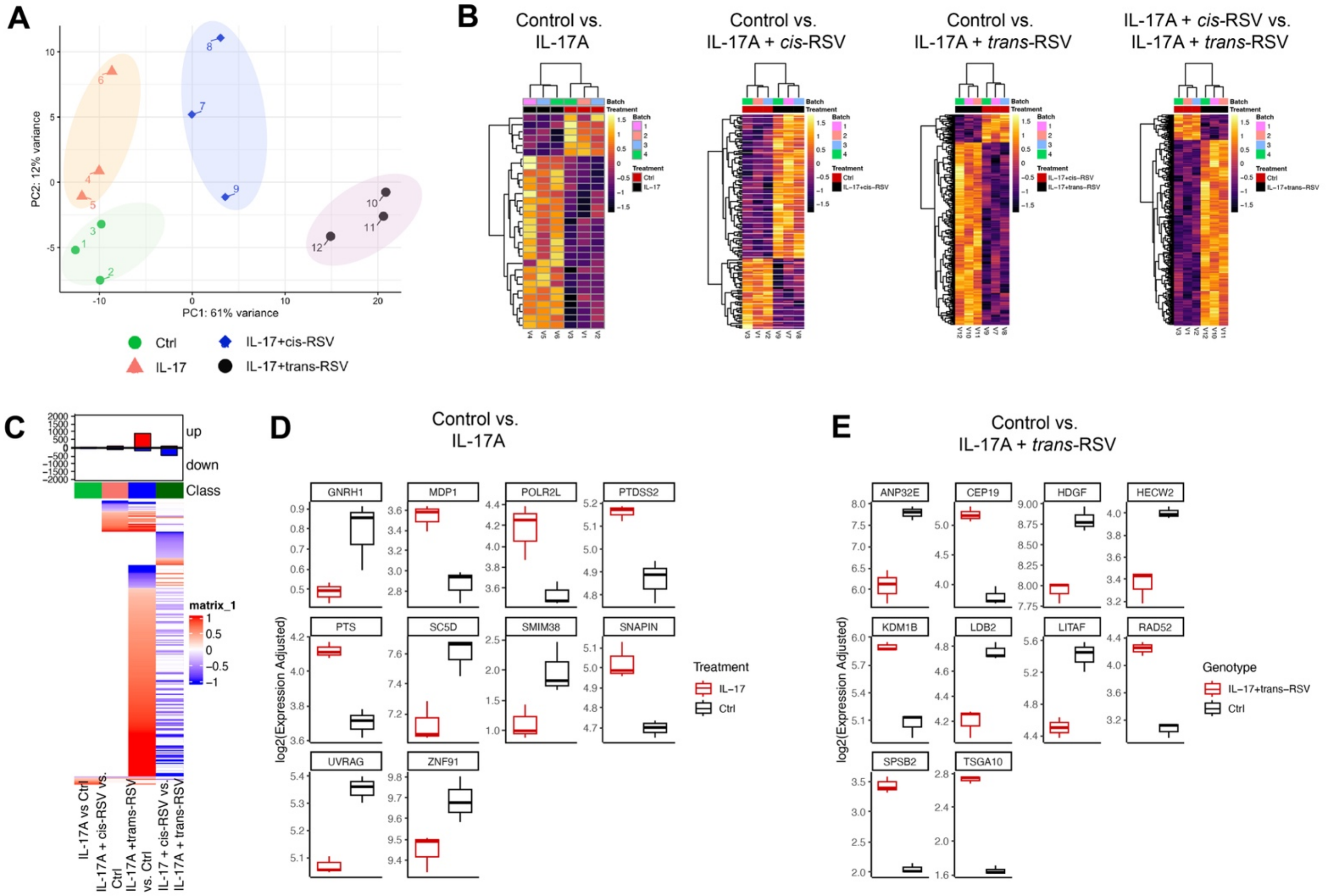


**Supplementary Figure S3. Transcriptome analysis of human neurons under different treatments.** (**A**) PCA plots show the individual experiments for each group, this includes control neurons (green), neurons treated with IL-17A (salmon), IL-17A + *cis*-RSV (blue), and IL-17A + *trans*-RSV (black). (**B**) Heatmaps show the top differentially expressed genes for IL-17A vs. control, IL-17A + *cis-RSV* vs. control, Il-17A + *trans*-RSV vs. control and for IL-17A + *cis*-RSV vs. Il-17A + *trans*-RSV. Upregulated genes are in lighter colors and downregulated genes are in darker hues. (**C**) Comparison of up and downregulated genes across the different comparisons listed in B. (**D**) Representative box plots show the top differentially expressed genes for IL-17A (red) vs. control (black) from RNAseq dataset. (**E**) Representative box plots show the top differentially expressed genes for IL-17A + *trans*-RSV (red) vs. control (black) from RNA-Seq dataset.

**Supplementary Figure S4 related to main Figure 4**

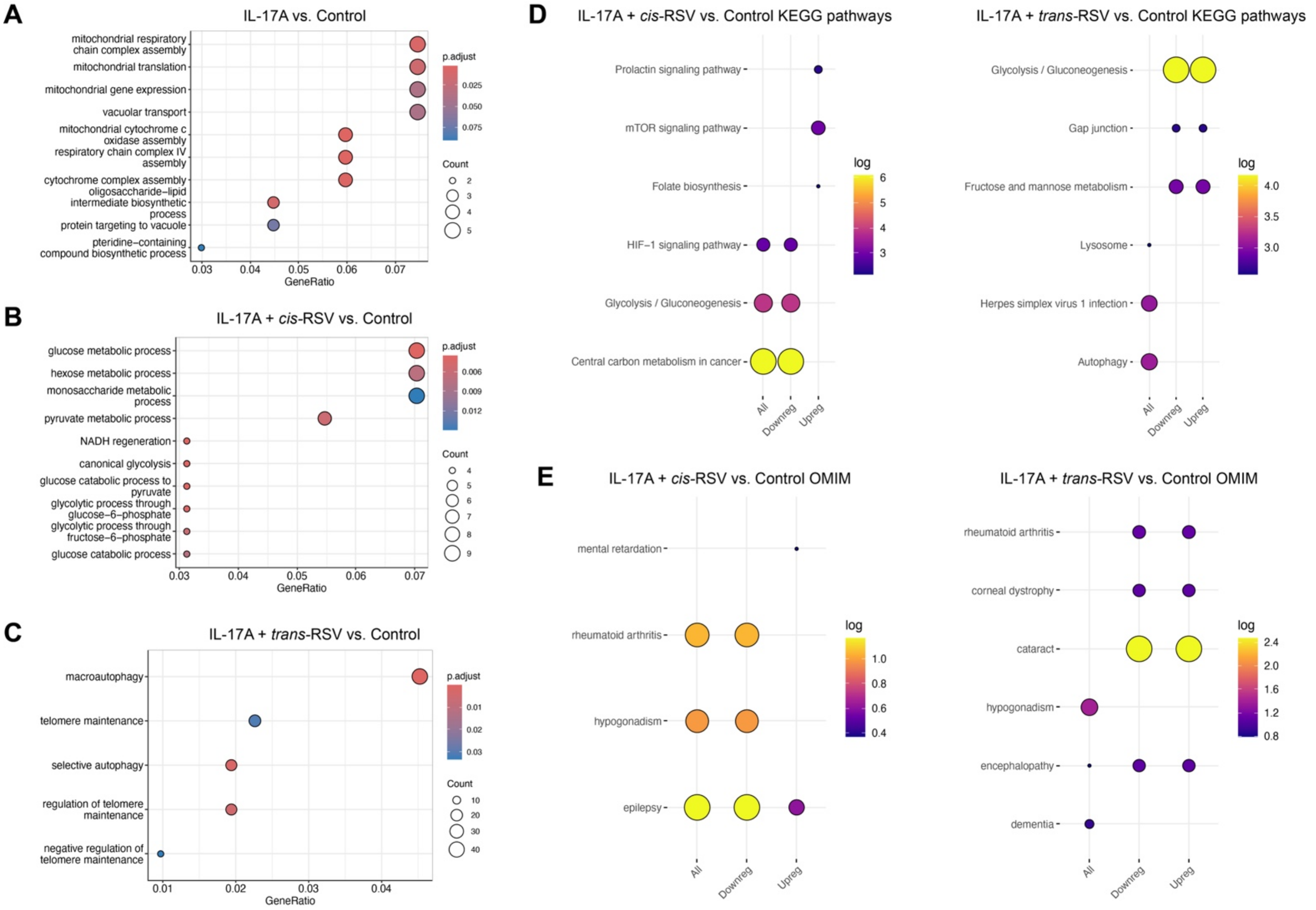


**Supplementary Figure S4. Functional enrichment analysis on human neurons under different treatments.** (**A-C**) Gene ontology analysis for molecular function shown for IL-17A vs. control (**A**), IL-17A + *cis-RSV* vs. control (**B**), and IL-17A + *trans*-RSV vs. control (**C**). (**D-E**) **Functional enrichment analysis using Enrich R** for either IL-17A + *cis-RSV* vs. control or IL-17A + *trans*-RSV vs. control for KEEG pathways (**D**), and for OMIM disease pathways (**E**).

**Supplementary Figure S5 related to main Figure 5**

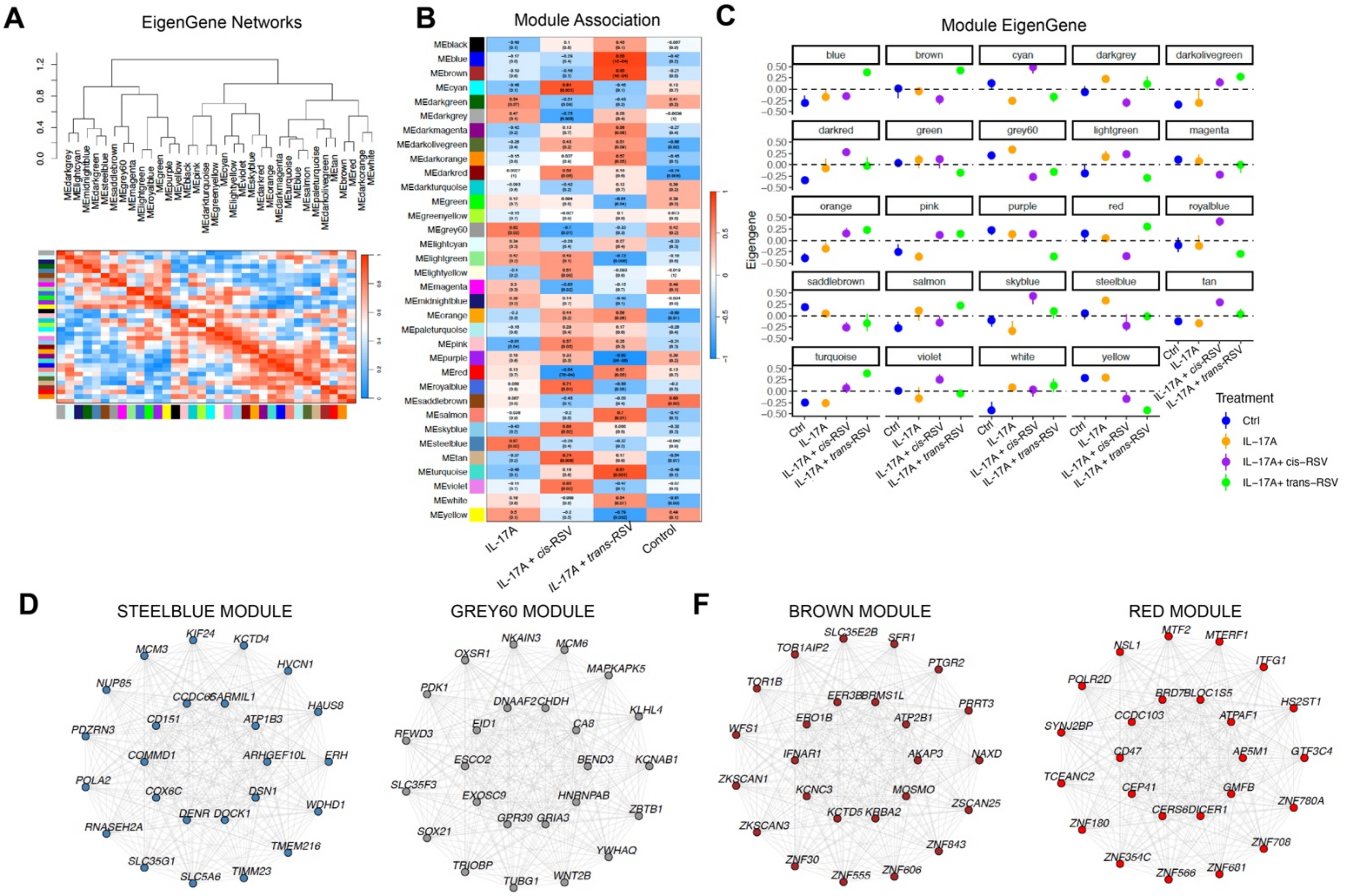


**Supplementary Figure S5. WGCNA analysis in relation to IL-17A, *cis* and *trans*-RSV in human neurons.** (**A**) Eigengene network analysis shows relationship between networks. (**B**) Heatmap shows the average change in gene networks co-expression per module. (**C**) WGCNA modules are shown for control (blue), IL-17A (yellow), IL-17A + *cis*-RSV (purple), and IL-17A + *trans*-RSV (green). (**D-F**) Representative gene networks are shown for steelblue and grey60 modules (**D**) and for brown and red modules (**F**).

**Supplementary Figure S6 related to main Figure 6**

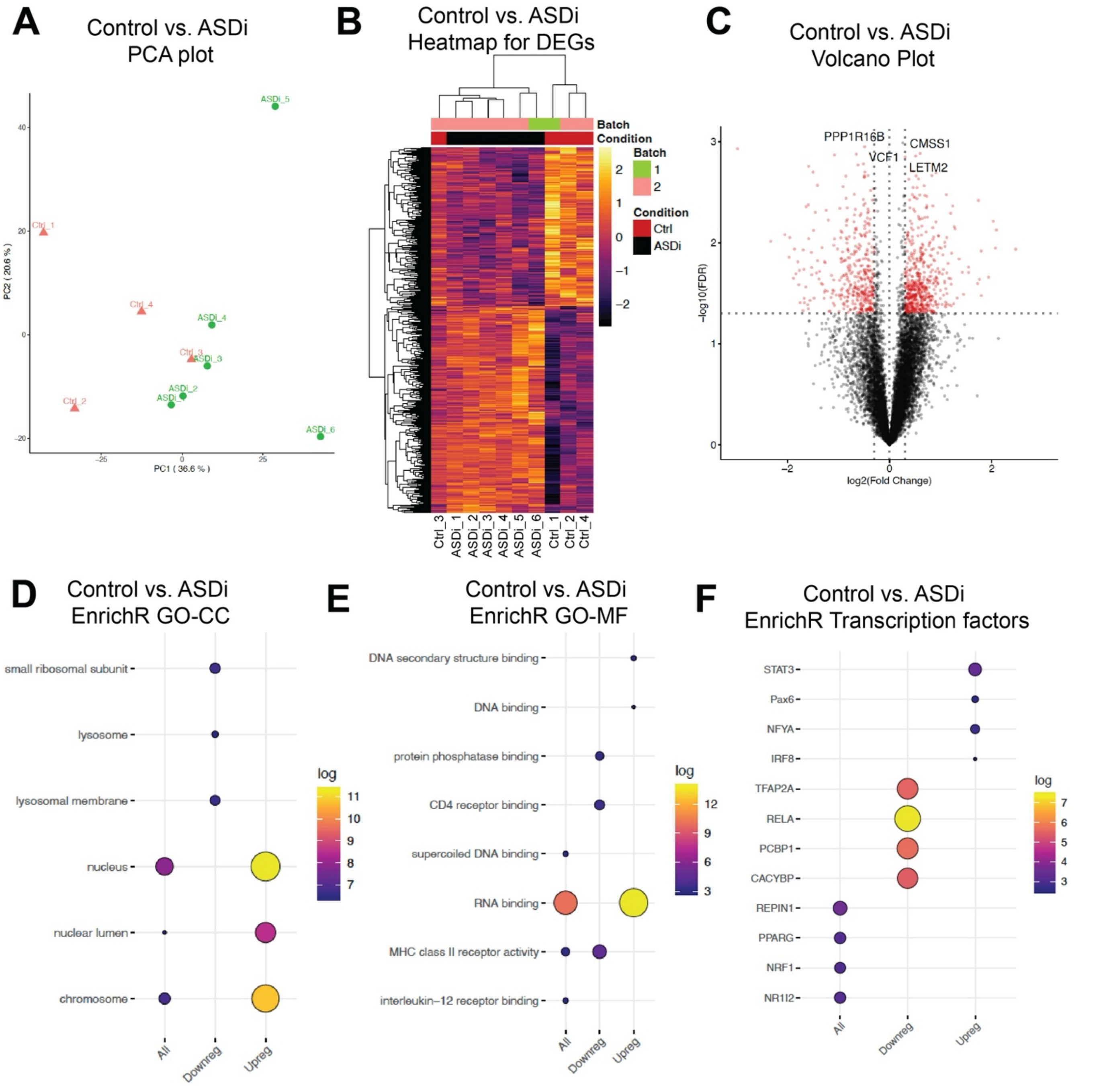


**Supplementary Figure S6. Transcriptome analysis of control and ASD patient lymphoblastoid cells.** (**A**) PCA plots show the individual lymphoblastoid control (red) and patient lines (green). (**B**) Heatmaps show the top differentially expressed genes for control (red) and ASD patient lines (black). Fold change is shown with upregulation in lighter colors and downregulation in darker hues. (**C**) Volcano plots showing up and downregulated DEGs raising above significance in red. (**D-F**) Functional enrichment analysis using Enrich R is shown for analysis of cellular compartment (CC) (**D**), molecular function (MF) (**E**) and for transcription factors (**F**).

### SUPPLEMENTARY TABLE LEGENDS

**Supplementary Table S1**
RNA seq analysis shown for human neurons treated with IL-17A and Resveratrol isomers

**Supplementary Table S2**
Functional enrichment analysis for human neurons treated with IL-17A and Resveratrol isomers

**Supplementary Table S3**
Gene ontology analysis for each WGCNA module is shown for IL-17A and Resveratrol isomers

**Supplementary Table S4**
EnrichR analysis for all WGCNA modules is shown for IL-17A and Resveratrol isomers

**Supplementary Table S5**
Clinical description of unaffected and ASD patients from whom lymphoblastoid cells were utilized for transcriptome studies

**Supplementary Table S6**
RNA seq analysis shown for lymphoblastoid cells from control and ASD individuals

**Supplementary Table S7**
Functional enrichment analysis is shown for lymphoblastoid cells from control and ASD individuals
